# Comparison of Illumina MiSeq and the Ion Torrent PGM and S5 platforms for whole-genome sequencing of picornaviruses and caliciviruses

**DOI:** 10.1101/705632

**Authors:** Rachel L. Marine, Laura C. Magaña, Christina J. Castro, Kun Zhao, Anna M. Montmayeur, Alexander Schmidt, Marta Diez-Valcarce, Terry Fei Fan Ng, Jan Vinjé, Cara C. Burns, W. Allan Nix, Paul A. Rota, M. Steven Oberste

## Abstract

Next-generation sequencing is a powerful tool for virological surveillance. While Illumina® and Ion Torrent® sequencing platforms are used extensively for generating viral RNA genome sequences, there is limited data comparing different platforms. We evaluated the Illumina MiSeq, Ion Torrent PGM and Ion Torrent S5 platforms using a panel of sixteen specimens containing picornaviruses and human caliciviruses (noroviruses and sapoviruses). The specimens were processed, using combinations of three library preparation and five sequencing kits, to assess the quality and completeness of assembled viral genomes, and an estimation of cost per sample to generate the data was calculated. The choice of library preparation kit and sequencing platform was found to impact the breadth of genome coverage and accuracy of consensus viral genomes. The Ion Torrent S5 outperformed the older Ion Torrent PGM platform in data quality and cost, and generated the highest proportion of reads for enterovirus D68 samples. However, indels at homopolymer regions impacted the accuracy of consensus genome sequences. For lower throughput sequencing runs (i.e., Ion Torrent 510 or Illumina MiSeq Nano V2), the cost per sample was lower on the MiSeq platform, whereas with higher throughput runs (Ion Torrent 530 or Illumina MiSeq V2) the cost per sample was comparable. These findings suggest that the Ion Torrent S5 and Illumina MiSeq platforms are both viable options for genomic sequencing of RNA viruses, each with specific advantages and tradeoffs.

## INTRODUCTION

Conventional Sanger sequencing has been the gold standard for genomic analysis of pathogens in public health laboratories for over three decades. However, the expansion of next-generation sequencing (NGS) technologies has increased demand for high-throughput sequencing of genomes at a lower cost (1). NGS has been used extensively for routine surveillance and outbreak investigation of numerous viral RNA pathogens. The exponential growth of genomic information generated for important pathogens has provided increased resolution for molecular epidemiology, as well as information necessary for the design of clinical assays and therapeutics (2–5). NGS methods are also useful for identifying pathogens in syndromes where etiologies often remain unknown (e.g., encephalitis, febrile illness), complementing or even replacing current diagnostic methods (2, 6, 7).

Over the past several years, the suppliers of high-capacity short-read sequencers have been reduced to two manufacturers: Illumina (sequencing-by-synthesis technology) and Thermo Fisher Scientific (Ion Torrent semi-conductor sequencing technology) (3). Illumina platforms have been used to generate nearly 90% of NGS data worldwide (https://www.wired.com/2016/02/gene-sequencing-goliath-wants-get-bigger-still/). Illumina produces several benchtop and production-scale sequencers with data outputs varying from 1.2 gigabases (Gb) to 6 terabases (Tb). In microbial research laboratories, the MiSeq platform is convenient for sequencing small microbial genomes (i.e., viruses and bacteria), compared to the larger-output Illumina platforms, that are more appropriate for eukaryotic genomes or very large studies, due to the balance of system/reagent costs and required sequencing depth (8–10). Similarly, the Ion Torrent technology is available in several models, producing data outputs from 30 megabases (Mb) to 25 Gb per chip. The Ion Torrent PGM, and newer systems (Ion Torrent S5, S5 XL, and GeneStudio S5, S5 Plus and S5 Prime) are also commonly used for microbial targeted-amplicon and whole-genome sequencing (8, 11–13).

Despite the extensive use of these platforms worldwide, there are limited studies providing a comprehensive comparison of yield and quality of generated data, as well as cost per sample to obtain complete viral RNA genomes. Comparing these NGS platforms is challenging due to their unique sequencing chemistries, resulting in vastly different quality score estimates and error profiles for the resulting data (14–16). Direct comparison of samples sequenced using both platforms is the ideal strategy to evaluate the advantages and limitations. Previous studies have mostly focused on 16S ribosomal genes or whole-genome sequencing of bacterial genomes on Sanger, Pacific BioSciences, 454 GS Junior, Ion Torrent, and Illumina platforms (8, 13, 17–19). In this study we sequenced a panel of 16 specimens known to contain enterovirus (EV) D68, poliovirus, norovirus, parechovirus and/or sapovirus using sequencing kits of varying output on the Illumina MiSeq, Ion Torrent PGM, and Ion Torrent S5 platforms.

## MATERIALS AND METHODS

### Sample Preparation

Sixteen samples were selected for the platform comparison: twelve clinical specimens, including nasopharyngeal (NP) swabs and stool specimens, and four cell culture isolates that were spotted on Whatman FTA cards. The chosen specimens contained picornaviruses (samples EV-D68-1 through −4 and Polio-5 through −8), caliciviruses (samples Noro-9 through - 12 and Sapo-15 and Sapo-16), or mixtures of both (samples Sapo-13; Parecho-13 and Sapo-14; Parecho-14) (Table S1). For NP swabs and stool specimens, samples were first clarified by centrifugation at 15,300 x g for 10 min. To remove host cellular debris and bacteria, 160 μl of the clarified supernatant was filtered through a sterile 0.45 μM Ultrafree-MC HV filter (EMD Millipore, Billerica, MA USA) by centrifugation at 3800 x g for 5 min at room temperature. Resulting filtrates were treated with Turbo DNase (Thermo Fisher Scientific, Carlsbad, CA USA), Baseline Zero DNase (Epicentre, Madison, WI USA), and RNase A (Roche, Pleasanton, CA USA) for 1 h at 37°C to degrade free nucleic acids. For all specimens, nucleic acids were extracted using the QIAamp Viral RNA Mini Kit (Qiagen, Germantown, MD USA) with optional on-column DNase treatment according to the manufacturer’s instructions (no carrier RNA) and eluted using 60 μl of Qiagen buffer AVE.

### Reverse Transcription and Random Amplification

Samples were processed using sequence-independent single-primer amplification (SISPA) (20, 21). First, viral RNA was reverse-transcribed using SuperScript IV reverse transcriptase (Thermo Fisher Scientific) and a 28-base primer consisting of a 3’ end with eight random nucleotides (N1_8N; CCTTGAAGGCGGACTGTGAGNNNNNNNN).

Second-strand extension was performed using Klenow 3’ → 5’ exo^−^ fragment (New England BioLabs, Ipswich, MA USA). Double-stranded cDNA was amplified using AmpliTaq Gold polymerase (Thermo Fisher Scientific) and N1 primer (CCTTGAAGGCGGACTGTGAG) under the following PCR conditions: 95°C for 5 min, 5 cycles of [95°C for 1 min, 59°C for 1 min, and 72°C for 1.5 min], followed by 25 cycles of [95°C for 30 sec, 59°C for 30 sec, and 72°C for 1.5 min with an incremental increase in the extension time of 2 sec per cycle]. Amplification was verified using the TapeStation 2200 (Agilent Technologies, Santa Clara, CA USA) prior to Agencourt AMPure XP bead purification (Beckman Coulter, Brea, CA USA; 1.8X ratio). Purified DNA was quantified using the Qubit dsDNA BR Assay kit (Thermo Fisher Scientific).

### Library Preparation and Sequencing

Sample dilution and library construction were performed with halved reactions according to the manufacturer’s instructions for the three library preparation kits evaluated: Nextera XT DNA Library Prep Kit (Illumina, San Diego, CA USA) and KAPA HyperPlus Kit (Roche) for Illumina sequencing, and the KAPA DNA Library Preparation Kit for Ion Torrent sequencing. Enzymatic shearing (included as part of the KAPA HyperPlus Kit) was not performed since cDNA fragments produced after SISPA are small enough for input directly into library construction. Individual barcoded libraries were visualized on the TapeStation 2200 before AMPure XP bead cleanup (1.8X ratio). Purified libraries were quantified prior to pooling using the LabChip GX (PerkinElmer, Waltham, MA USA) for Nextera XT libraries and KAPA libraries sequenced on the Ion Torrent S5, whereas KAPA HyperPlus libraries and libraries sequenced on the Ion Torrent PGM platform were quantified by qPCR using the NEBNext Library Quant Kit for Illumina (New England BioLabs) or the KAPA Library Quantification Kit for Ion Torrent platforms (Figure 1). Multiplex Illumina libraries were sequenced by using MiSeq 500v2 and Nano 500v2 kits (2 x 250 basepair (bp) paired-end runs). The Ion Torrent PGM libraries were prepared using the IC 200 kit for Ion Chef (Thermo Fisher Scientific) and sequenced on the Ion Torrent PGM using the 316 and 318 semi-conductor sequencing chips, while the Ion Torrent S5 libraries were prepared using the “Ion 510™ & Ion 520™ & Ion 530™” for Ion Chef Kit for 400 base-read libraries and sequenced on the Ion Torrent S5 using an Ion 510 semi-conductor sequencing chip (Thermo Fisher). For reporting of results and discussion, the eight dataset names are abbreviated as follows: PD6 and PD8 for library preparation with the KAPA DNA Kit and sequencing on an Ion Torrent PGM 316 v2 chip and 318 v2 chip, respectively; MKN and MK5 for library preparation with the Kapa HyperPlus Kit and sequencing on an Illumina Nano 500 v2 run and Illumina 500 v2 run, respectively; MNN and MN5 for library preparation with the Nextera XT Kit and sequencing on an Illumina Nano 500 v2 run and Illumina 500 v2 run, respectively; and SDG and SDS for library preparation with the KAPA DNA Kit and sequencing on an Ion Torrent S5 510 chip. The S5 datasets are distinguished by whether the libraries were size-selected using E-Gel SizeSelect II gels (SDG dataset, 300 bp; Invitrogen, Carlsbad, CA USA) or purified using standard AMPure XP bead cleanup (SDS) prior to quantification and chip loading (Figure 1).

**Figure 1.**
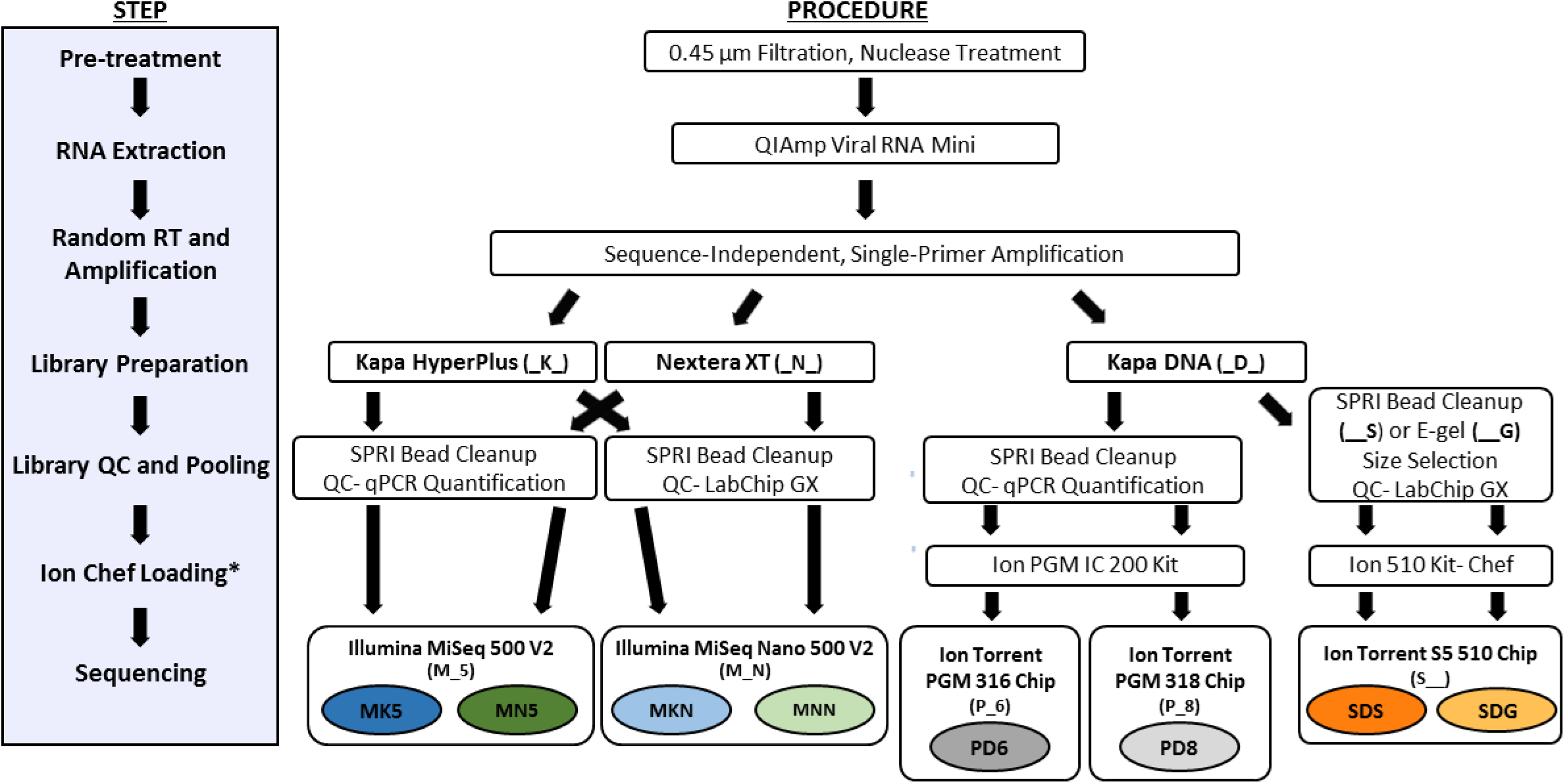
Overview of library preparation and sequencing kits utilized for preparing viral specimens for sequencing on the Illumina, Ion Torrent PGM and Ion Torrent S5 platforms. Abbreviations for each dataset based on the type of library kit and sequencing kit/cartridge used: NexteraXT 500v2 (MK5), NexteraXT Nano 500v2 (MNN), KAPA HyperPlus 500v2 (MK5), KAPA HyperPlus Nano 500v2 (MKN), KAPA DNA Ion Torrent 316v2 (PD6), KAPA DNA Ion Torrent 318v2 (PD8), KAPA DNA Ion Torrent S5 510 SPRI Size Selection (SDS), KAPA DNA Ion Torrent. 510 E-Gel Size Selection (SDG). *Ion Chef loading is only performed for Ion Torrent sequencing runs.

### Viral Genome Analysis

Sequencing data were processed using a custom viral bioinformatics pipeline (VPipe), accessible to partner public health researchers through the CDC SAMS partner portal (https://sams.cdc.gov/). Human reads were identified and removed through read mapping to the human genome (h19) using bowtie2 (22). Adaptors, primer sequences, and low-quality bases (phred score threshold of 20) were trimmed from the raw reads, followed by removal of duplicate reads. Filtered datasets were assembled using SPAdes v.3.7 (23) with multiple kmer lengths and settings specific for either Illumina or Ion Torrent datasets. Resulting contigs were compared to the NCBI non-redundant nucleotide database and an in-house database of viral sequences using blastn and blastx (24). Geneious v.11.1.2 (25) (BioMatters, Newark, NJ USA) was used to map sequencing reads to their respective contigs, using the map-to-reference tool with sensitivity set to low/fastest with a fine tuning of three iterations. Reference recruitments were manually evaluated for accuracy and trimmed to produce the final consensus sequence generated by *de novo* assembly. For each sample, consensus genomes from all eight datasets were aligned to generate the longest consensus sequence. This “master” consensus provided a consistent reference for performing a second reference-based recruitment for calculating the proportion of target reads and coverage statistics. For samples with fewer target reads (EV-D68-1 through 4, and Sapo-16) the closest genome in GenBank was used as the master consensus (Table S2). The filtered fastq files for all datasets have been submitted to the NCBI SRA database (BioProject PRJNA550105).

### Statistics

To assess differences in the proportion of sequences removed during quality control filtering between samples/datasets, a generalized linear model was fitted with the SAS proc glimmix procedure (SAS Institute, Cary, NC). Beta distribution was utilized with logit link function because read proportion is a percentage variable (26). The response variable was fitted on observed variables “virus”, “dataset”, and “library kit”. Variable “dataset” is nested within variable “library kit” since each dataset (produced on a given sequencing technology) can be only used with a specific compatible library preparation protocol (variable “library kit”). Least-square means were calculated using Tukey comparisons to account for multiple comparisons across different scenarios (27). To compare genome coverage across datasets, Pearson’s correlation coefficient was computed using JMP statistical software (version 9.0.0; SAS, Cary, NC, USA) (28). EV-D68 datasets were not considered for the correlation analysis due to low coverage across multiple datasets.

### Cost Analysis Calculation

The cost per sample was calculated for sequencing preparation workflows performed in this study, plus an estimate of the cost per sample for sequencing on an Ion Torrent S5 530 chip (which has higher sequencing data output than the S5 510 chips used in this study). The pricing of all kits and consumables utilized from pretreatment and extraction through sequencing was included, taking into account the total number of samples which could be processed by a given kit and the multiplexing level for the sequencing run considered. For consistency, the LabChip GX HS assay was used for calculating the cost of library quantitation for all preparations, despite using both LabChip GX and qPCR-based quantitation methods for this study. Sample and reagent shipment, equipment, and personnel costs were not considered.

## RESULTS

### Sequencing Yield

The eight datasets analyzed were sequenced using five different chips/kits which vary in their advertised read output (Figure 1, Table S3): Ion Torrent PGM 316 v2 chip (PD6), Ion Torrent PGM 318 v2 chip (PD8), Ion Torrent S5 510 chip (SDS, SDG), Illumina MiSeq 500v2 Nano kit (MKN, MNN), and standard Illumina MiSeq 500v2 kit (MK5, MN5). Total sequencing yield per run (Table S4) was within the output ranges claimed by manufacturers, with two exceptions. For the Ion Torrent PGM runs (PD6 and PD8), where the total yield was roughly a third of that expected, decreased yields were likely due to less efficient chip loading and lower proportions of clonal and useable reads with the PGM platform relative to the newer S5 platform (Table S5). Lower yields were also observed for Illumina libraries prepared using the KAPA HyperPlus Kit (MKN, MK5) compared to the Nextera XT kit (MNN, MN5). This was attributed to lower clustering densities on the Illumina MiSeq (MKN, 478K/mm^2^ and MK5, 439K/mm^2^ vs. MNN, 1120K/mm^2^ and MN5, 1046K/mm^2^), despite using qPCR for library quantitation, which is thought to provide more accurate estimates of sample concentration than electrophoresis-based methods (29).

### Data Yields after Quality Control

For all libraries, prefiltering of raw fastq files consisted of removal of host (human) sequences, trimming of low quality bases and adapters, and removal of short (<50 bp) and duplicate reads. After quality control, 17.3-46.1% of total reads were retained per library (Table S4). The proportion of reads removed during each step of the quality control filtering varied greatly by virus and sample (Figure 2). A large proportion of host reads (56.5-98.4%) were removed for EV-D68 samples (NP swabs), regardless of the library preparation kit and sequencing platform used (Figure 2A, Table S6, p<0.0001). There was also a significant difference in the proportion of host reads removed for stool specimens (samples Noro-9 through Sapo-16) compared to cell culture specimens (samples Polio-5 through Polio-8). The greatest loss of data for cell culture and stool specimens was due to removal of duplicate sequences (Figure 2B-D), except in the case of samples sequenced on the Ion Torrent PGM platform (PD6, PD8), where removal of low quality/short reads led to the greatest loss of data (Table S7, p<0.0001). The proportion of duplicate reads removed was greater for samples sequenced on standard Illumina 500 v2 runs (MK5, MN5) compared to Illumina Nano 500 v2 runs (MKN, MNN) and Ion Torrent S5 runs (SDS, SDG) (Table S8, p<0.0001).

**Figure 2.**
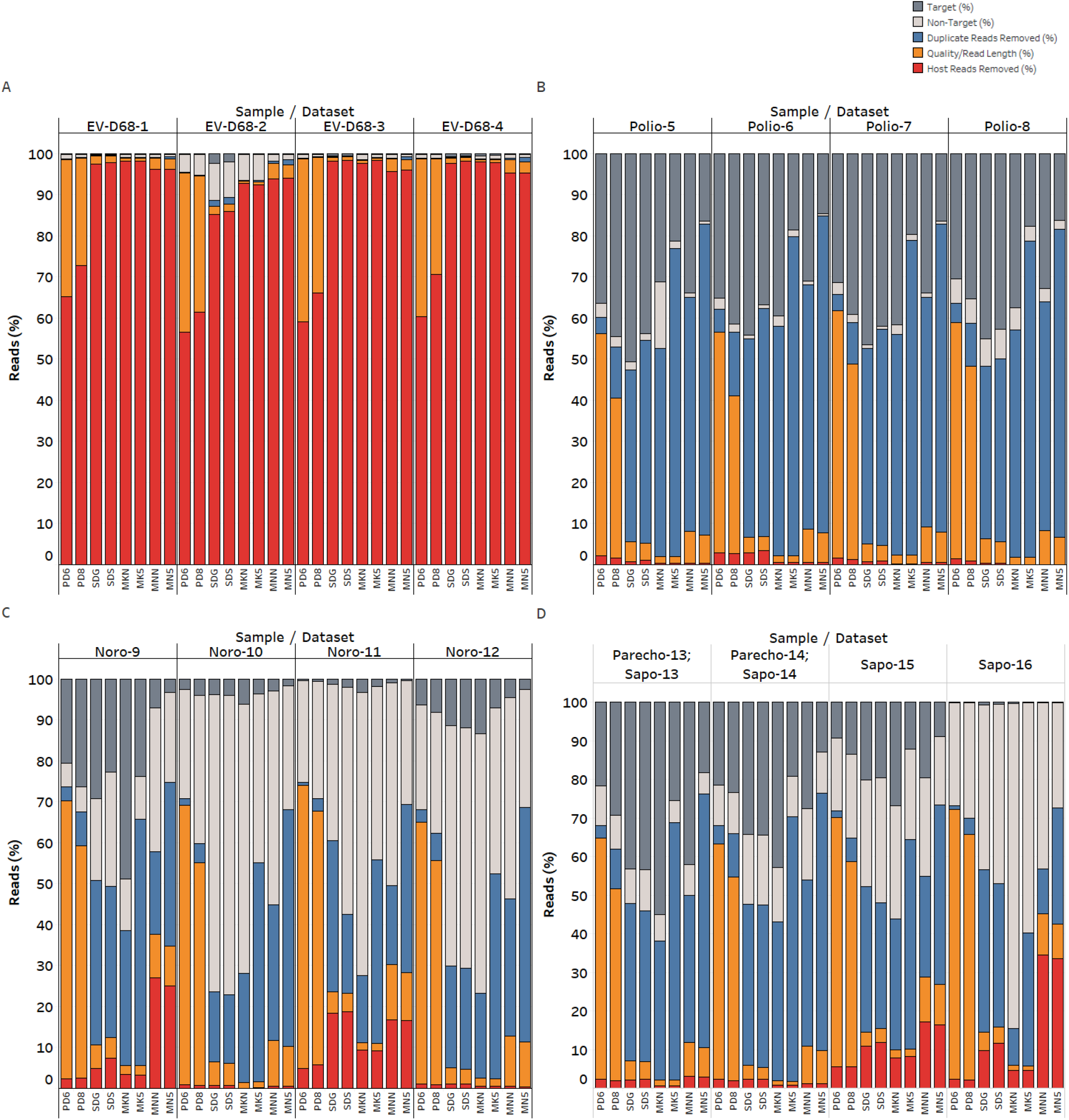
Results of fastq quality filtering for each sample/dataset. Samples are separated by target virus: EV-D68 1-4 (Panel A), polio 5-8 (Panel B), norovirus 9-12 (Panel C), and sapovirus/parechovirus 13-14 and sapovirus 15-16 (Panel D). The top label on the x-axis indicates the sample, while the bottom x-axis label indicates the NGS dataset. Each stacked bar represents the total reads per dataset. The percentage of reads removed at each filtering step is denoted by color, including the percentage of host/human reads removed (red), the proportion of sequences removed which were less than 50 bp after quality and adapter trimming (orange), and the proportion of duplicate reads removed (blue). Reads remaining after filtering are indicated by the gray bars, with the light gray bars corresponding to non-target (i.e., non-viral) sequences and the dark gray bars corresponding to target viral sequences.

Because of the increase in read duplication with sequencing depth, the proportion of viral (i.e., target) reads did not scale linearly with sequencing output. Rather, datasets with intermediate sequencing output (MKN, SDG and SDS) tended to have a higher proportion of viral reads per sample (Figure 3A). Regardless of whether duplicate reads were considered, the greatest proportion of viral reads were observed for polio samples (Figure 3B), whereas low sequencing yields were obtained for EV-D68 samples despite the high titer of virus measured in the original specimens (Ct values of 17 to 21.6 using an EV-D68-specific qPCR assay, Table S1). Illumina datasets prepared using the Kapa HyperPlus Kit (MKN, MK5) and datasets generated using the Ion Torrent S5 platform (SDG, SDS) consistently produced the highest proportion of target reads for norovirus and EV-D68 samples, respectively (Figures 3A and 3B). For norovirus samples, where specimens comprised a larger span of Ct values (from 18 to 27 using a norovirus-specific qPCR assay), a general trend of decreasing target reads with increasing Ct was observed (Figure S1). However, when comparing EV-D68 and sapovirus samples, which had a narrower distribution of Ct value, there was no obvious correlation between Ct and the amount of target sequence data obtained (Figure S1). For example, only 0.1-0.6% of reads mapped to Sapo-16 (Figure 3), which had a relatively low Ct value of 18.9.

**Figure 3.**
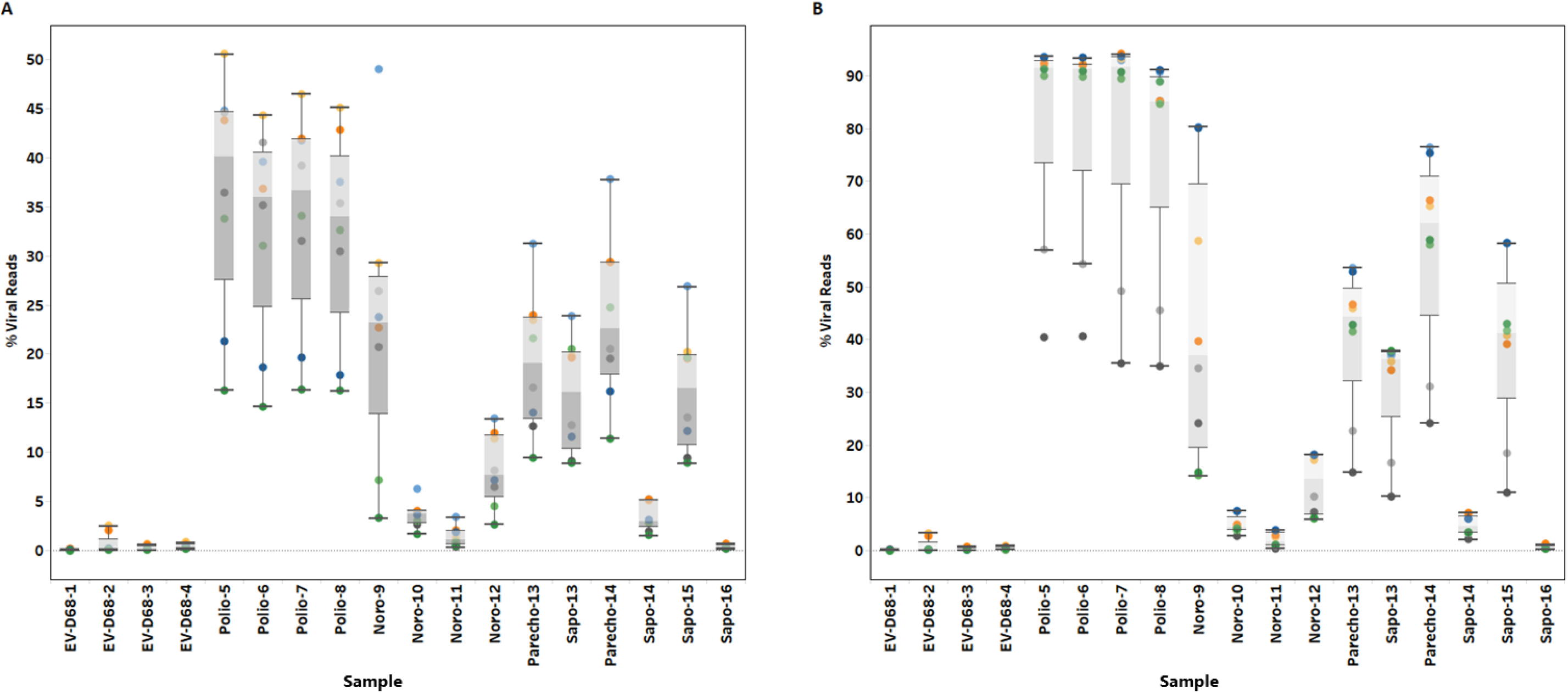
The effect of library preparation and sequencing strategy on the proportion of viral (target) reads obtained for a given sample. Each point represents the percent viral reads for a given dataset, denoted by color. Box-and-whisker plots depict the range of percent viral reads for each sample. Whiskers extend to 1.5 times the interquartile range. The grey zones indicates the upper and lower quartiles, and the line between the two quartiles indicates the median percent target reads. Panel A depicts the analysis of the percentage of viral reads after all quality control filtering steps (see Methods), whereas in Panel B, duplicate reads were considered in the analysis.

### Comparison of Genome Coverage

When trying to generate genome sequences, the breadth of coverage (i.e., percentage of positions in a genome which are sequenced), as well as the depth of coverage (i.e., number of reads covering a given position in the genome) influence the completeness and accuracy of genome sequences produced (30). Considering the breadth of coverage across target viruses (Figure 4), at ≥1X read coverage the Ion Torrent S5 datasets (SDG, SDS) generated the most consistent coverage for EV-D68 genomes, while the MK5 dataset produced the greatest breadth of coverage for norovirus samples. Ion Torrent S5 and Illumina MiSeq datasets all performed well for sequencing of poliovirus; for parechovirus samples, the breadth of genome coverage was within 10 bp of the master consensus length for all datasets. If only genome positions with ≥10X read coverage were considered for calculating the breadth of coverage, the MK5 dataset covered the greatest proportion of the genome for 14 of the 18 viruses sequenced (Figure 4).

**Figure 4.**
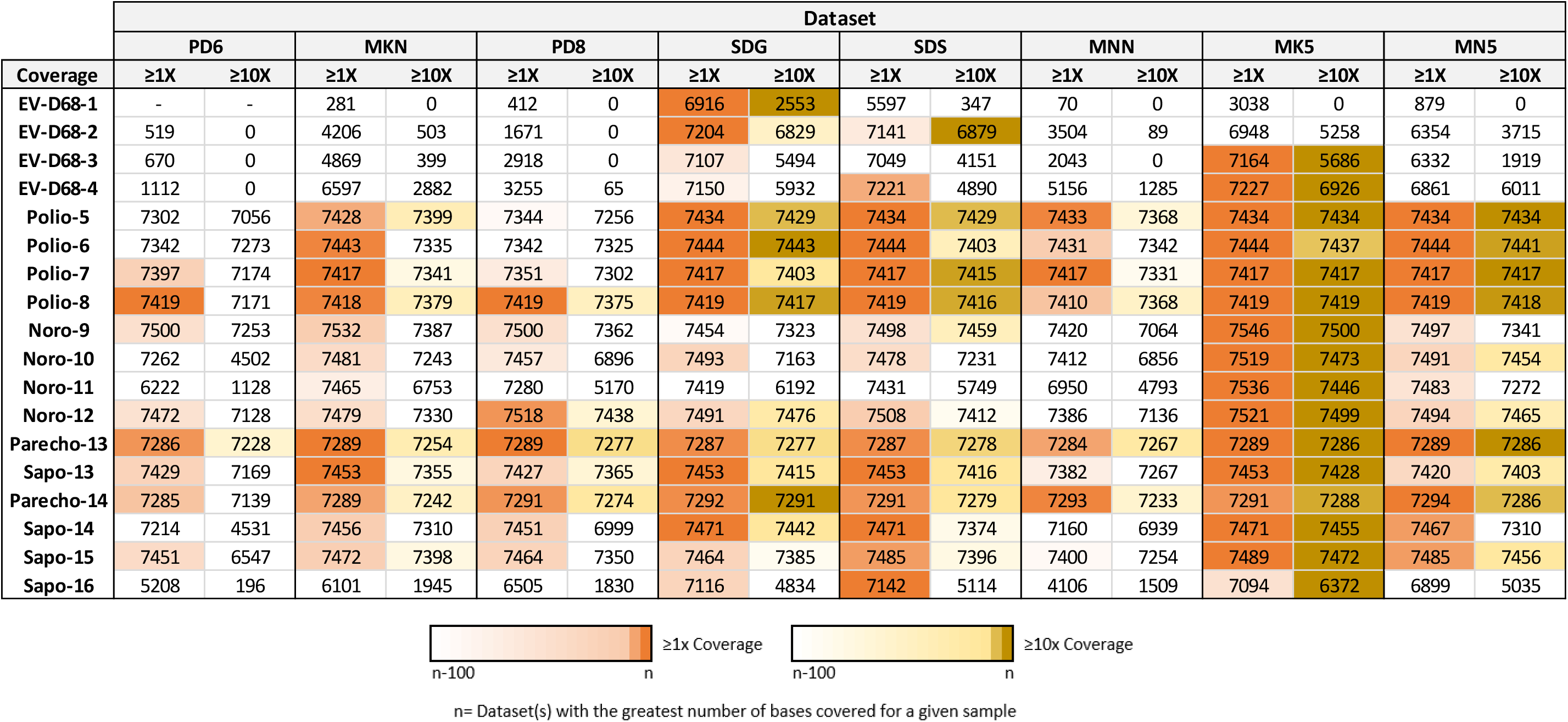
Breadth of coverage across target genomes. Heatmap indicating the total number of bases (genome positions) for each sample which had at least 1X read coverage and 10X read coverage per dataset. Cells highlighted in orange (for ≥1X coverage) and yellow (for ≥10X coverage) indicate datasets that were within 100 bp of the dataset with the greatest number of bases covered. Datasets with the greatest coverage for a given sample correspond to cells with the darkest color.

Considering the pattern of sequencing coverage across a genome, reproducible peaks in the coverage profiles were observed, as shown for poliovirus samples for example (Figure 5). Despite uneven coverage profiles produced by the SISPA protocol (31–33), a relatively small number of reads (compared to bacterial or eukaryotic genomes) was needed to reconstruct near-complete genomes (approximately 30,000 reads to obtain at least single read coverage across >99% of the genome, or ≥10X read coverage across >98% of the genome, for viruses with ~7.3-7.5 kb genomes, Figures S2 and S3). While all datasets compared produced statistically similar coverage patterns, libraries prepared using the same library preparation kit had a stronger correlation, particularly for MiSeq libraries prepared using the Nextera XT kits (MNN and MN5) and Kapa HyperPlus kit (MKN and MK5) (Dataset S1, p<0.0001). For Ion Torrent PGM datasets, PD6 coverage patterns were consistently most similar to PD8. Interestingly, PD8 datasets were also very similar to SDS datasets, with PD8 datasets demonstrating the strongest correlation to SDS datasets for 10 of 14 viruses with sufficient coverage for comparison (Supplemental Dataset S1). The E-gel size selection (prior to library pooling) may have influenced the final distribution of fragment sizes, leading to differences in the coverage patterns between SDG and SDS datasets.

**Figure 5.**
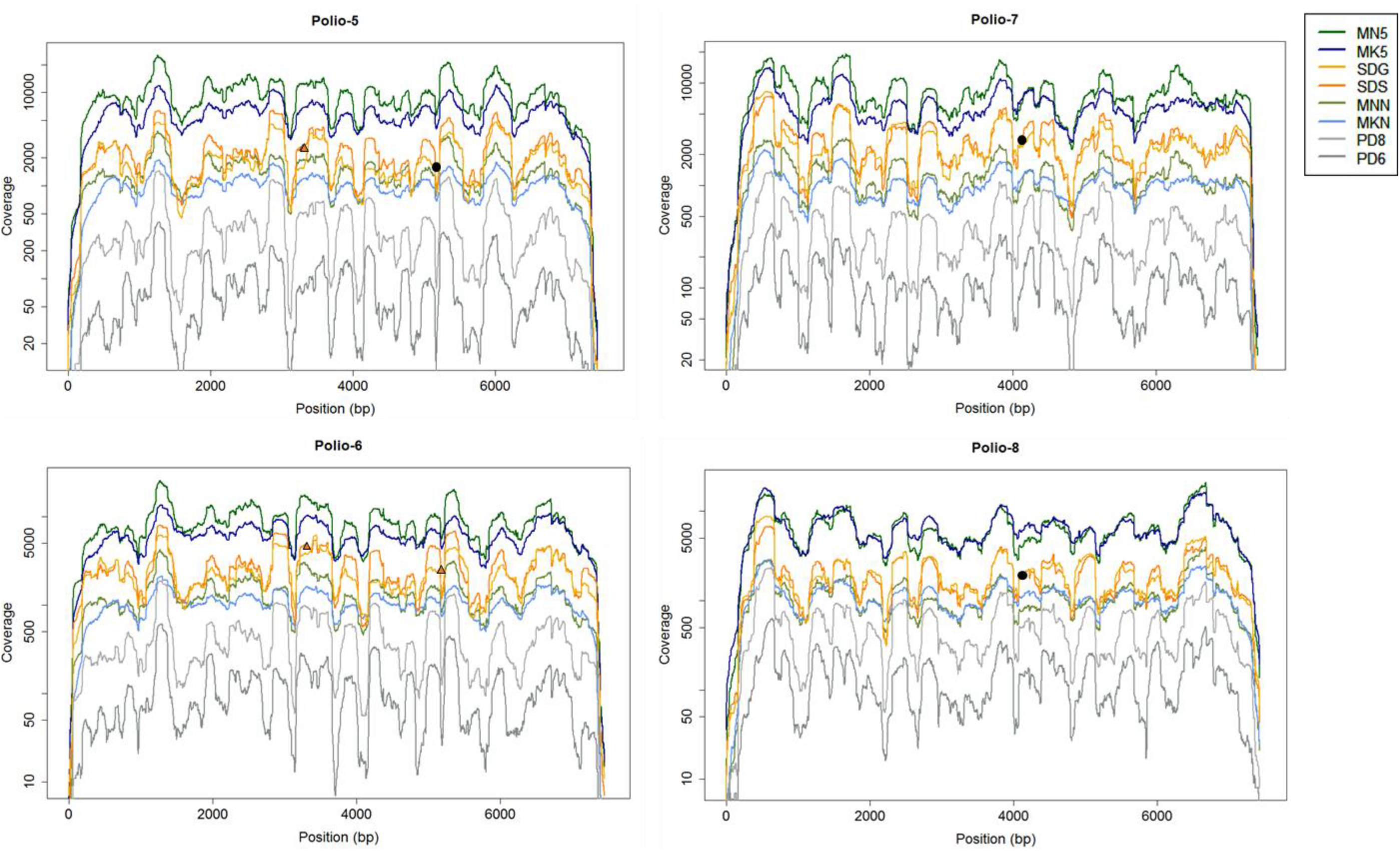
Coverage patterns across the poliovirus genome. The depth of coverage, plotted on a log scale, across the length of the genome is depicted for all datasets (denoted by color). Polio-5 and Polio-6 are both type 1 polioviruses, while Polio-7 and Polio-8 are type 3 viruses. Orange triangles indicate the positions of high frequency indels in the SDS consensus genome sequences, while black points indicate the positions of high-frequency indels found at the same position for both SDG and SDS datasets (only one point per position is shown for simplicity).

### Accuracy of Viral Consensus Genome Sequences

Indels were observed in genome consensus sequences generated from Ion Torrent datasets, even in areas with high read coverage. Indels (insertions) in Ion Torrent S5 datasets were observed in two locations for Polio-5 and Polio-6 samples, and one location for Polio-7 and Polio-8 samples (Figure 5). These locations correspond to homopolymer runs of seven or eight C residues for poliovirus type 1, and a homopolymer run of six A residues for poliovirus type 3 (Table S9). At some positions, an indel was observed in only one of the two Ion Torrent S5 datasets (SDS or SDG). In these scenarios, the indel frequency was still high for both datasets, but only one exceeded the 50% threshold where an indel would be called in the final majority consensus. Indels in consensus sequences were also observed in Ion Torrent datasets for norovirus, parechovirus, and sapovirus samples (Table S9). While indels for SDS and SDG sequences were always single-nucleotide insertions at areas of homopolymer repeats, indels detected in PD6 and PD8 consensus sequences did not always occur at repeat regions and were often deletions rather than insertions.

### Cost Analysis

The calculated cost per sample decreased substantially with increased levels of multiplexing, particularly at moderate levels of multiplexing (Figure 6). As multiplexing levels were increased, the cost per sample reached a plateau, since certain reagent costs will always scale linearly with the number of samples processed. This includes the cost of pretreatment, reverse transcription, library preparation, and nucleic acid quantitation/quality control consumables (Table S10). The total cost per sample when sequencing 16 samples on an Illumina MiSeq 500V2 Nano run was $76.25 and $81.07 using the Nextera XT and Kapa HyperPlus kits, respectively, compared to $129.38 and $134.20 when sequencing on a standard Illumina MiSeq 500V2 run. The cost per sample for an Ion Torrent S5 510 chip run closely matched the cost per sample of an Ion Torrent PGM 318v2 run ($124.18 and $125.04 respectively when sequencing 16 samples, Figure 6), with the S5 510 chip producing more high quality reads with a shorter run time than the PGM 318 chip (Figure 2, Table S4) (34). When comparing the Ion Torrent S5 and the Illumina MiSeq system, the difference in the cost per sample decreases with increased multiplexing. For example, when sequencing only one sample, the difference in cost per sample between an Ion Torrent S5 530 run and an Illumina MiSeq 500v2 run (MK5 preparation), which have roughly comparable read outputs, is $65.88 ($1352.08 vs $1286.20), compared to $5.47 ($113.97 vs $108.50) when multiplexing 24 samples. For lower read output runs (i.e., Ion Torrent S5 510 vs Illumina MiSeq 500v2 Nano), the cost per sample is markedly lower for the Illumina MiSeq 500v2 Nano (Figure 6).

**Figure 6.**
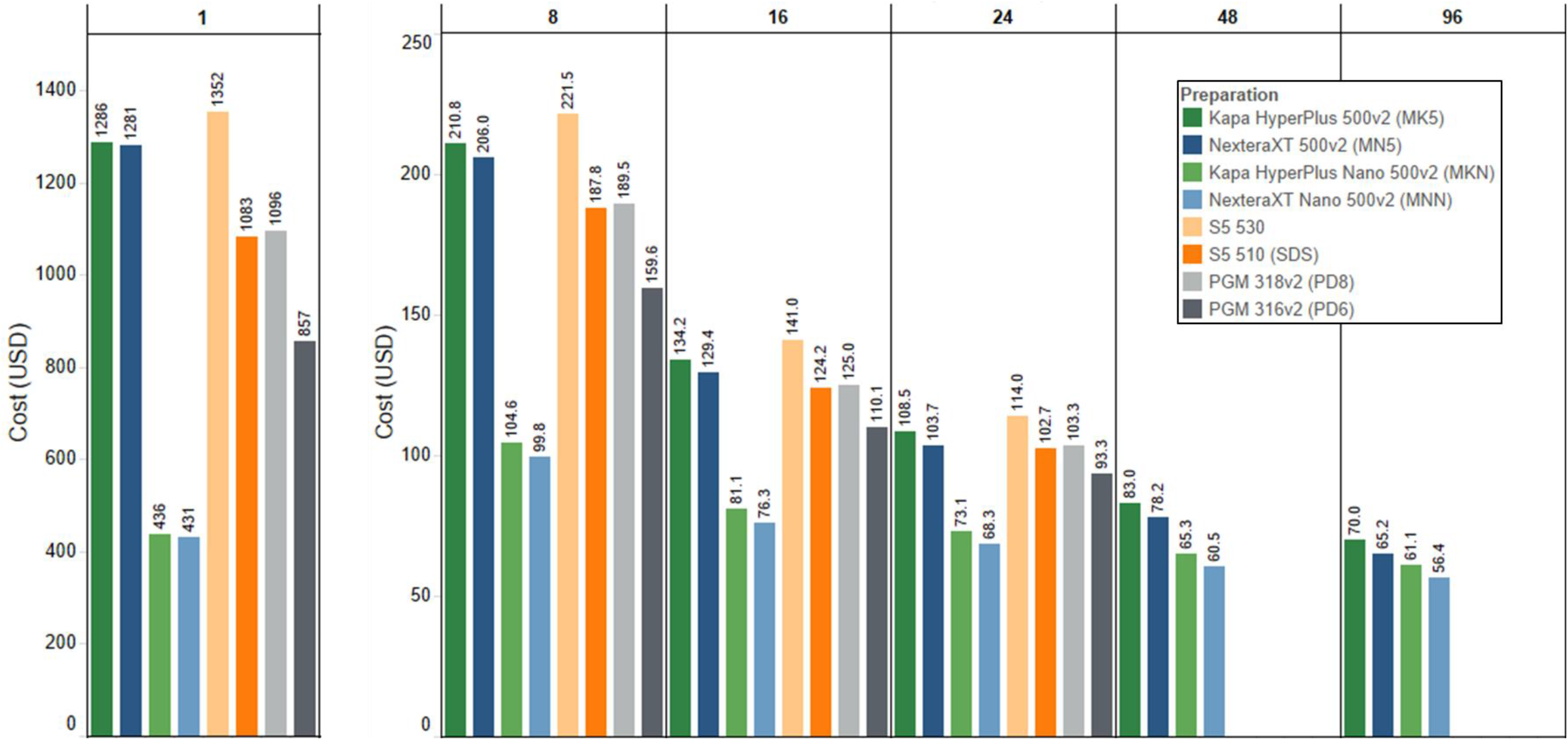
Estimated cost per sample for performing next-generation sequencing based on kits used for sequencing and the level of multiplexing. From left to right, each block represents the number of samples multiplexed in a single run. Individual bars correspond to the library preparation and sequencing kit used. The number above each bar indicates the estimated cost per sample. The Ion PGM and S5 calculations are only performed out to multiplexing levels of 24 samples, as the KAPA DNA library kit currently only makes 24 unique indices. Calculations include the cost of reagents, kits and consumables from sample pretreatment through sequencing (Fig. 1).

## DISCUSSION

Sixteen samples containing RNA viruses were multiplexed and sequenced using eight different combinations of library preparation and sequencing kits to evaluate the ability of each strategy to produce target viral genomes. Datasets with intermediate output (MKN, SDS, and SDG) were found to have the highest proportion of viral reads. While the number of target reads increased with the amount of data generated, the removal of a greater proportion of duplicate reads led to lower proportions of target reads in Illumina MiSeq 500 v2 runs (MK5, MN5). A similar finding was reported in a study optimizing methodologies for sequencing of human respiratory syncytial virus, with higher proportions of duplicate reads observed in the higher output Illumina NextSeq 500 datasets compared to the MiSeq (35). This is most likely due to over-amplification of viral genomes during SISPA, combined with a greater probability with increasing sequencing depth of generating duplicate reads by chance, especially for small genomes (36). Even when duplicate reads are retained, differences in the proportion of target reads were observed between datasets. Libraries prepared using the Kapa HyperPrep kit consistently had the highest proportion of target reads for norovirus samples, while Ion Torrent S5 libraries consistently produced relatively more data for EV-D68 samples. For the Kapa HyperPrep libraries, the lower proportion of reads removed during the host removal and quality filtering stages may have contributed to higher yields of target reads. In addition, better breadth and depth of coverage was observed for samples prepared with the KAPA library kits compared to the Nextera XT kit. This was particularly prominent for caliciviruses, where even KAPA datasets with lower total read output had better breadth of genome coverage than Nextera XT datasets (e.g., MKN, SDG, and SDS datasets vs. MNN, and MK5 vs MN5). The required tagmentation/fragmentation step in the Nextera XT protocol likely leads to a greater loss of coverage over genome termini due to sequence selection bias (37–39).

Indels were observed in eight consensus genomes for the Ion Torrent S5 datasets, and six consensus genomes for the Ion Torrent PGM datasets. It is well documented that the predominant base-call error produced by Ion Torrent semiconductor sequencing platforms is indels, particularly after long homopolomeric stretches (8, 16, 17, 40). Interestingly though, high-frequency indels observed in the PGM datasets (PD6, PD8) were almost always deletions rather than insertions, and were not typically associated with homopolymer repeats, in contrast to S5 datasets. A previous study examining error bias in Ion Torrent PGM data identified single-base high-frequency indel errors which were not associated with long homopolymer repeats and were unique to a single run (14). This observation is similar to the patterns observed in our Ion Torrent PGM datasets, where the location of high-frequency indels manifesting in genome consensus sequences were usually only observed in one of the two PGM datasets. The disparity in the location and nature of high frequency indels between the Ion Torrent PGM and S5 platforms suggests that there may be differences in the flow-value accuracy and resultant error profiles for these two Ion Torrent devices. While indels can be corrected for viruses that are well-characterized, particularly for the S5 dataset where indels were only observed in regions of homopolymer repeats of the same nucleotide, they may pose a challenge for genome sequencing of novel or relatively uncharacterized viruses.

When designing NGS experiments, the choice of multiplexing level and sequencing kit (i.e., the depth of sequencing per sample) will depend on the anticipated proportion of non-target (e.g., bacterial, human) reads relative to target, and the total number of samples which ultimately need to be sequenced for a given experiment. For example, poliovirus and other enteroviruses are known to shut down host RNA transcription early in infection, thus increasing the proportion of viral RNA relative to host RNA in virus isolates (41). Therefore, a greater number of enterovirus isolates can be multiplexed in one run— greater than 96 on a standard Illumina MiSeq or Ion Torrent S5 530 run for experiments with a large number of samples, or 24 samples on an Illumina MiSeq Nano or Ion Torrent S5 510 run for smaller experiments (21). Conversely, clinical samples have more variability in the proportion of target reads even when sequencing samples with similar qPCR Ct values. Additional factors such as the specimen type, the age of the specimen, the proportion of non-target nucleic acids (e.g. in a respiratory or fecal sample), and the stability of the pathogen being targeted likely influence whether complete genomes are obtained. For metagenomic sequencing directly from patient specimens such as stool, it is advisable to limit sequencing runs to 16-24 samples on a standard MiSeq or Ion Torrent 530/540 run. Even lower multiplexing levels (or sequencing kits with greater output) would be necessary for sequencing of EV-D68 from nasal swabs. In these situations, a targeted NGS method, such as generating EV-D68 amplicons prior to library preparation and sequencing, is likely the most cost-effective option (42, 43). Ideally, researchers should strive to sequence as many samples as possible on a run, as multiplexing dramatically decreases the cost per sample.

Researchers may also decrease the cost through reducing library preparation reaction volumes, as this is typically the most costly step in NGS preparation (Table S10). While reducing reaction volumes deviates from the formulations validated by manufacturers, many researchers (including ourselves) have used half-reactions for preparing NGS libraries with no noticeable effect on quality, and other studies have reported reliable library preparation down to one-sixteenth reactions (44–47).

Our study has several limitations. While the reported results are broadly applicable to laboratories that sequence RNA viruses, only a subset of RNA viruses (picornaviruses and caliciviruses) were evaluated in this study. SISPA was used for random reverse transcription for all datasets which likely influenced the pattern of genome coverage to a greater degree than the library preparation or sequencing platform used. Despite the documented biases of SISPA, this method is still commonly used for RNA viruses, especially for samples where enrichment of RNA is necessary to obtain enough starting material for library construction (48). We also did not evaluate any targeted NGS methods, which are likely more effective when performing routine sequencing for particular viral pathogens (49). Nevertheless, this study complements previous research investigating the utility of Ion Torrent and Illumina platforms (8, 13, 17–19, 50–54). As more public health laboratories begin to implement NGS, these results provide important considerations in weighing the advantages and disadvantages of using a particular sequencing platform or library preparation kit for performing metagenomic sequencing of RNA viruses.

## Acknowledgements

This work was made possible by Federal appropriations to the Centers for Disease Control and Prevention (CDC) through the Advanced Molecular Detection (AMD) line item. This research was also supported in part by appointments to the Research Participation Program at CDC administered by the Oak Ridge Institute for Science and Education (L.C.M., C.J.C., and M.D-V.) through an interagency agreement between CDC and the U.S Department of Energy.

We would like to thank Nikail Collins for her assistance with preparation of norovirus samples used in this study.

## Appendix A. Supplementary data

Supplementary tables and figures: Supplement_PlatformCompare_JCM.pdf Supplementary data set: DatasetS1_SupplementalStatistics.xlsx

